# Species comparison of liver proteomes reveals links to naked mole-rat longevity and human aging

**DOI:** 10.1101/220343

**Authors:** Ivonne Heinze, Martin Bens, Enrico Calzia, Susanne Holtze, Oleksandr Dakhovnik, Arne Sahm, Joanna M. Kirkpatrick, Karol Szafranski, Natalie Romanov, Kerstin Holzer, Stephan Singer, Maria Ermolaeva, Matthias Platzer, Thomas Hildebrandt, Alessandro Ori

## Abstract

Mammals display wide range of variation in their lifespan. Investigating the molecular networks that distinguish long-from short-lived species has proven useful to identify determinants of longevity. Here, we compared the liver of long-lived naked mole-rats (NMRs) and the phylogenetically closely related, shorter-lived, guinea pigs using an integrated omic approach. We found that NMRs livers display a unique expression pattern of mitochondrial proteins that result in distinct metabolic features of their mitochondria. For instance, we observed a generally reduced respiration rate associated with lower protein levels of respiratory chain components, particularly complex I, and increased capacity to utilize fatty acids. Interestingly, we show that the same molecular networks are affected during aging in both NMR and humans, supporting a direct link to the extraordinary longevity of both species. Finally, we identified a novel longevity pathway and validated it experimentally in the nematode *C. elegans*.

## Introduction

Among mammals lifespan generally correlates with other life-history parameters such as gestation period and body mass [1]. In this perspective, a subterranean rodent, the naked mole-rat (*Heterocephalus glaber,* NMR), and humans represent two species outliers by having an exceptionally long lifespan relatively to their body mass. NMRs are eusocial animals that live in colonies where only a subgroup of animals is devoted to reproduction (usually a queen and one male called pasha) [2]. NMRs exhibit other exceptional traits including lifelong fertility, resistance to infection, high regenerative capacity, resistance to cancer and diabetes, reviewed in,[3,4]. For these reasons, NMRs have drawn attention of multiple studies aimed at identifying the molecular mechanisms behind their extreme longevity and resistance to age-related diseases. Comparative genome analysis has revealed positively selected genes in NMR [5,6], and RNA-seq analysis revealed minimal changes in gene expression during aging [5,7], supporting the view of enhanced maintenance of homeostasis in NMRs at the molecular level. NMRs possess enhanced protein stability and increased proteasomal activity [8,9], negligible levels of cellular senescence [10], over-activation of pathways that contribute to stress resistance (e.g., the nuclear factor erythroid2-related factor 2 (NFE2L2, previously NRF2/Nrf2) and p53) [3,11], atypical expression of extracellular matrix components, such as high molecular mass hyaluronan, that confer resistance against cancer development [12]. Intriguingly, NMRs have higher steady-state levels of oxidative damage compared to, e.g., mouse [8,13], and possess mitochondria with unusual morphology in the heart and skeletal muscle [14]. However, NMRs appear to be protected from the age-dependent increase in oxidative damage that manifest in other species [8], presumably due to enhanced detoxifying systems [15].

There is a wealth of evidence linking energy metabolism to the aging process and organism lifespan. Dietary interventions and nutrient sensing pathways have been shown to play a central role in modulating aging in different organisms [16–18]. Studies of genes under positive selection pressure across related species that show different lifespan have highlighted genes involved in mitochondrial homeostasis balance [19], likely influencing both energy metabolism and hormetic responses affecting lifespan [20]. Age-dependent changes of mitochondrial ultrastructure and activity have been described both in flies and mice [21].

There is emerging evidence that metabolic changes in the NMR contribute to adaptation to its ecosystem [22], allow to act as a “superorganism” with its eusocial life style [4], and might be related to its extreme longevity. However, the relationship between NMR metabolism and longevity has not yet been investigated. Given its central role in organism metabolism, we set out to investigate the liver of NMRs in order to identify novel molecular signatures of longevity. Since ecological adaptations are more likely to affect gene expression (Fraser 2013), and mechanisms of aging act both at the transcript and protein level [23], we performed a cross-species comparison between NMR and the shorter-lived guinea pig (*Cavia porcellus,* GP) using an integrated proteomic and transcriptomic approach. In order to investigate cross-species differences in the context of aging, we additionally analyzed livers from young and old NMRs, and related the identified changes to human aging by studying the liver proteome of 12 individuals aged between 31 and 88 years. Finally, we validated one of the newly identified longevity pathways to be a mediator of lifespan in the nematode *C. elegans*.

## Results

### Cross-species comparison of liver proteomes reveals elevated expression of detoxifying enzymes in naked mole rats

We first compared the liver proteomes of 4 young adult male NMRs (2.7-3.8 year old (yo)) and GPs (0.7-1 yo) using mass spectrometry (Table S1). For each animal, we obtained a quantitative proteome profile by liquid chromatography tandem mass spectrometry, and estimated absolute protein abundances using the iBAQ method [24]. In order to directly compare the two species, we mapped both NMR and GP proteins to the respective human orthologs, and used these as the reference for comparison (see Material and Methods). This allowed us to perform a quantitative comparison of the two species using estimated absolute abundances for 3248 protein groups quantified by at least two unique proteotypic peptides in both species (Figure 1A and Table S2). In order to validate our approach, we obtained RNA-seq data from the same samples and determined transcript-level fold changes between the two species. For this, only those reads that exclusively mapped to conserved regions were used, a method used for transcriptomic cross-species comparisons [7] (Table S2). Protein and transcript fold changes displayed a significant positive correlation (Pearson R=0.52, p<2.2e-16, considering all the cross-quantified cases; Pearson R=0.78, p<2.2e-16, considering all the cases significant in both dataset (adj. p <0.1); Figure 1B), which is in line with comparisons performed within the same species [25]. These data indicate that our strategy can reveal meaningful differences in protein abundance between species and that many of these changes are driven by changes in transcript levels.

**Figure 1.**
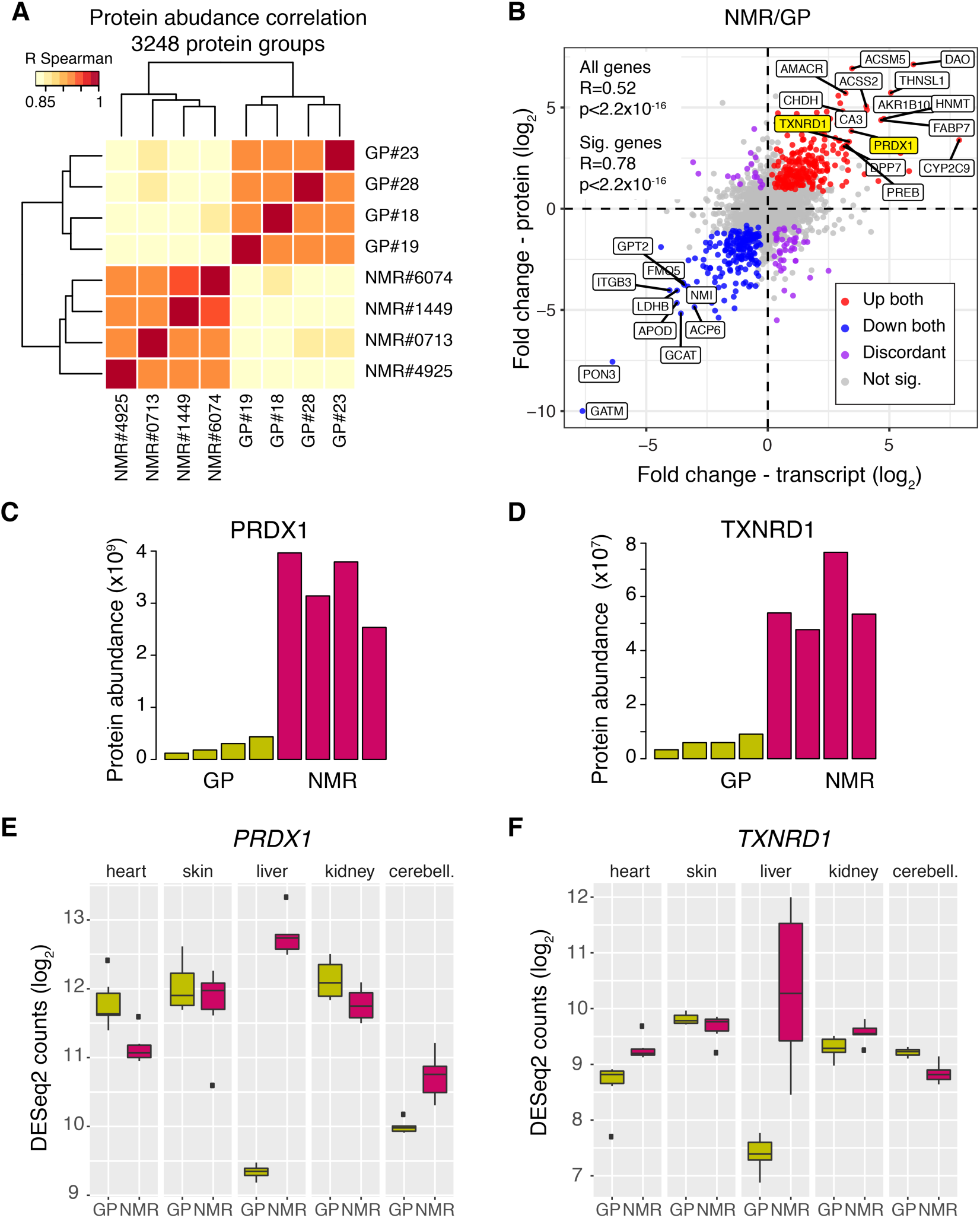
Cross-species comparison of the liver proteome integrated with RNA-seq from the same animals. **(A)** Liver proteomes from 4 adult naked mole rats (NMRs) and 4 guinea pigs (GPs) were compared using quantitative mass spectrometry. Hierarchical clustering based on the correlation between proteome profiles based on 3248 protein groups quantified across the two species. **(B)** Correlation between fold changes estimated at the protein level by quantitative mass spectrometry and at the transcript level by RNA-seq. Colored dots indicate genes significant (adj. p<0.1) in both comparisons. The names of selected genes that show consisted fold changes in the two comparisons are indicated. For RNA-seq analysis only reads mapping to conserved regions between the two species were considered. **(C and D)** PRDX1 and TXNRD1 as examples of longevity-associated proteins that show drastically increased abundance in NMR vs. GP. Each bar represents the abundance estimated for one animal. **(E and F)** Comparison of transcript levels of *PRDX1* and *TXNRD1* across multiple tissues. For both genes, transcript levels are increased in NMR vs. GP in the liver (q<2.2×10^-300^ for *PRDX1*; q=1.5×10^-45^ for *TXNRD1*), while they show similar abundances in the other tissues examined. Related to Tables S1 and S2.

We next asked whether our comparison would reveal known longevity-associated proteins. We found strikingly higher levels of peroxiredoxin 1 (PRDX1) and thioredoxin reductase 1 (TXNRD1) in NMRs as compared to GPs (Figure 1C and D). *PRDX1* and *TXNRD1* are both target genes of the transcription factor NFE2L2, they play a crucial role in maintaining cell redox homeostasis, and both have been shown to influence lifespan in multiple species by buffering ROS and promoting proteostasis [26–28]. Interestingly, both PRDX1 and TXNRD1 are cytosolic enzymes, and their mitochondrial counterparts (PRDX5, TXNRD2) are instead expressed at similar levels in both species (Table S2).

We next wondered whether we could identify similar differences in other organs. For this purpose, we compared RNA-seq data from heart, skin, kidney and cerebellum across the two species and found that increased transcript levels of *PRDX1* occur exclusively in the liver (Figure 1E and F). This suggests that increased level PRDX1 and TXNRD1 might be linked to a specific metabolic activity of the NMR liver.

### The liver of naked mole rats has a unique metabolism characterized by reduced mitochondrial respiration and enhanced lipid metabolism

Intrigued by this finding, we used gene set enrichment analysis to investigate differences in pathways and molecular networks between the long-lived and shorter-lived species. Our analysis returned pathways linked to energy metabolism (Figure 2A and Table S3). In particular, we found pathways related to lipid metabolism to be up regulated, and gene sets related to oxidative phosphorylation to be down regulated in NMR (q<0.05). Among up-regulated proteins involved in lipid metabolism, we found enzymes responsible for fatty acid beta-oxidation (e.g., ACOX2 and ACOX3), and lipid (e.g., ACACA and ACSL5), cholesterol (e.g., MVD and DHCR24) and bile acids biosynthesis (e.g., AMACR) compared to GP (Figure 2A and B and Table S2). Many of these are direct target of the nuclear receptor peroxisome proliferator-activated receptor alpha (PPARα), a master regulator of energy metabolism linked to aging [29] (Figure 2B). The majority of these enzymes showed higher abundance in the liver of NMRs both at transcript and protein level, irrespectively of their sub-cellular localization (Figure 2B).

**Figure 2.**
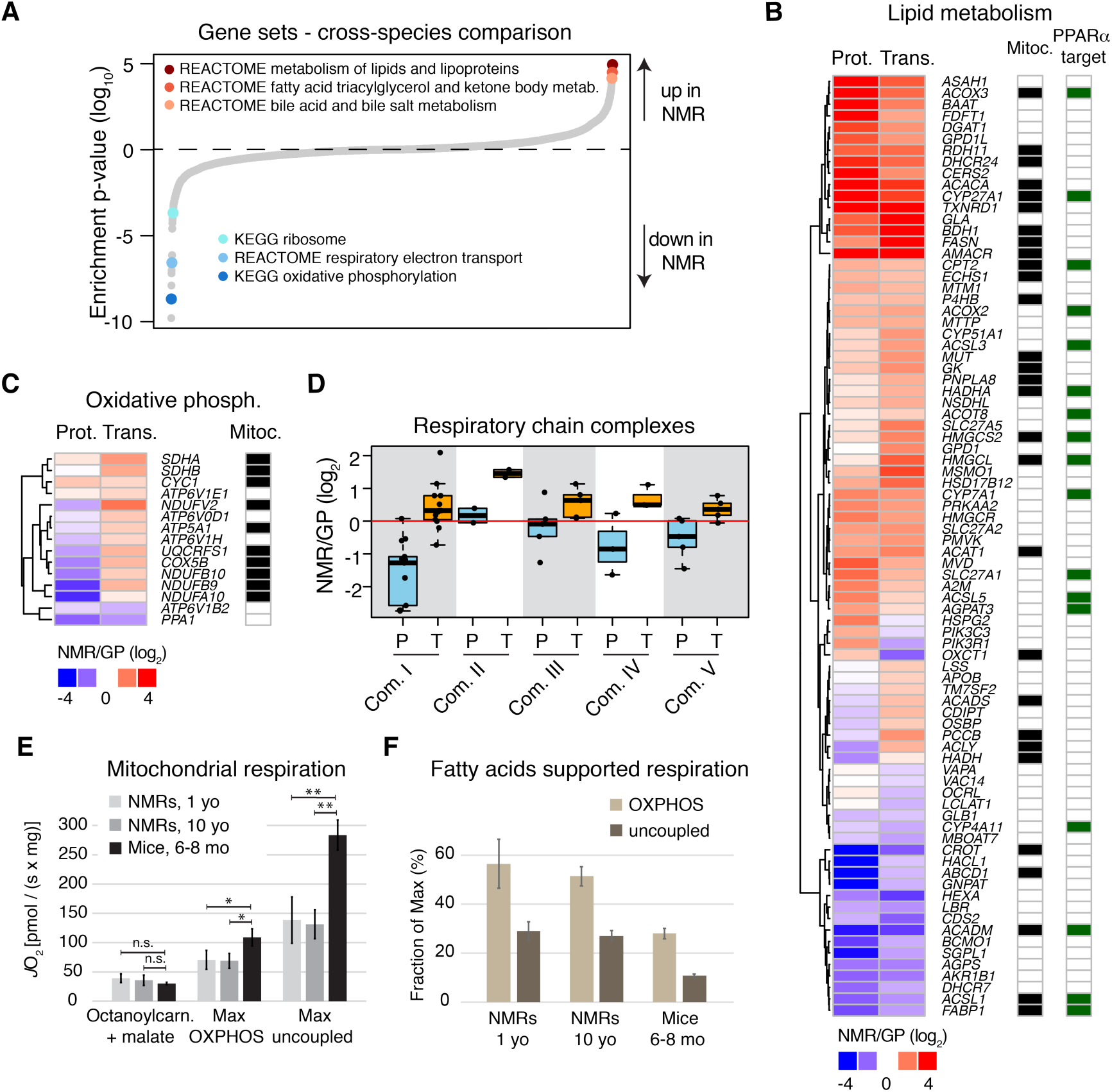
Distinctive features of NMR liver metabolism. **(A)** Gene set enrichment analysis was performed on proteomic data. Gene sets are plotted according to the log_10_ value of the calculated enrichment score. Positive and negative values are used for gene sets showing higher and lower abundance in NMR, respectively. Because of the redundancy among the significantly affected gene sets (q<0.05), we highlighted representative categories of gene sets. The complete list of enriched gene sets is available in Table S3. **(B)** Protein (P) and transcript (T) fold changes for genes involved in lipid metabolism (“REACTOME METABOLISM OF LIPIDS AND LIPOPROTEINS”, combined RNA-seq and proteome q<0.001), and **(C)** oxidative phosphorylation that are significantly affected in NMR vs. GP (selection criteria as in B). **(D)** Fold changes comparison for genes of the different complexes of the respiratory chain. Light blue and orange boxes indicate protein and transcript fold changes, respectively. **(E)** Mitochondrial oxygen flux supported by octanoylcarnitine and malate, compared to maximum coupled (OXPHOS) and maximum uncoupled respiration in liver of NMR (1 and 10 yo) and 6-8 mo male mice. **(F)** Ratio of fatty acid supported respiration to maximum OXPHOS and uncoupled respiration in liver of NMR and mouse. In E and F, reported values are averages obtained from n=4 animals per experimental group ± standard deviation. *=p<0.05; **=p<0.001; n.s=not significant. Related to Table S3.

In order to exclude that such differences would arise from a different organelle composition of the NMR liver cells, we analyzed the distribution of fold changes for proteins belonging to different cell compartments and found slight, but significantly lower general levels of mitochondrial proteins (mean log_2_ fold change=-0.17, p=0.01, Welch two sample t-test), and higher levels of lysosomal proteins (mean log_2_ fold change=0.2, p=0.005, Welch two sample t-test) in NMR compared to GP.

Regarding oxidative phosphorylation, NMRs showed reduced abundance of a subset of mitochondrial respiratory chain components (Figure 2C). Interestingly, these differences manifested almost exclusively at the protein level, and affected to different extents the respiratory chain complexes, with components of complex I being the most strongly reduced (mean log_2_ fold change=-1.53, p=7.2×10^-5^, Welch two sample t-test, Figure 2D). The opposite changes of mitochondrial enzymes involved in lipid metabolism and respiratory chain components indicate that NMR livers possess a distinct composition of their mitochondrial proteome.

In order to investigate whether proteomic differences result in altered mitochondrial activity of NMR liver parenchyma, we performed *ex vivo* measurements of cellular respiration in liver extracts of NMR by means of high-resolution respirometry (see Material and Methods). We measured mitochondrial respiration from an independent group of young (1 yo, n=4) and middle aged (10 yo, n=4) male NMRs and compared them to adult male mice (6-8 mo, n=4). We first compared the contribution of fatty acids to mitochondrial activity using octanoylcarnitine and malate as substrates. After this, maximum coupled respiration-state (OXPHOS-state) was established by further addition of glutamate and succinate. Finally, we completed the test sequence by adding carbonyl cyanide p-(trifluoromethoxy)-phenylhydrazone (FCCP) in order to achieve the maximum uncoupled respiration-state. We observed (Figure 2E) that NMRs and mice showed similar rates of mitochondrial respiration supported by fatty acids. In contrast, NMR livers show reduced maximum mitochondrial activity compared to mouse liver. In line with the reduced abundance of respiratory chain components, mitochondrial oxygen consumption normalized per wet tissue volume was significantly reduced in both maximum OXPHOS (35% (p=0.011) and 37% (p=0.014) in 1 and 10 yo NMRs, respectively; one-way ANOVA followed by Bonferroni’s t-test), and maximum uncoupled states (51% (p<0.001) and 54% (p<0.001)). This implies a ∼2-fold higher fatty acid supported mitochondrial respiration in NMRs compared to mice (Figure 2F). Taken together, our data indicate a marked rearrangement of energy metabolism in the liver of NMR characterized by an enhanced lipid metabolism and globally reduced level of mitochondrial respiration.

### Cross-species- and aging-related changes correlate

Given the existing evidences linking both lipid metabolism and oxidative phosphorylation to lifespan [30,31], we investigated how these pathways are affected during aging in NMRs. We therefore compared the liver proteome of young (3-4 yo, n=3) and old (>20 yo, n=3) NMRs using quantitative mass spectrometry (Table S1 and S4). Because of the rarity of aged NMR samples, the old NMRs were obtained from a different source (zoo) of the young ones. In order to exclude potential biases due to different origin of the two sample groups, we additionally compared two groups of young (1 yo, n=4) and middle-aged (>10 yo, n=4) NMRs sourced from the same facility (Table S1 and S4). Histological evaluation of NMR liver tissue derived from the different age groups did not reveal significant inflammatory, fatty or fibrotic changes, while an increased lipofuscin pigment deposition (a known aging marker) was detectable with aging (Figure S1A).

Unsupervised hierarchical clustering based on the obtained proteome profiles confirmed an effect of aging on the NMR liver proteome in both dataset (Figure 3A and S1B). In addition, protein fold changes positively correlated across the two dataset (Pearson R=0.23, p<2.2×10^-16^, considering all the cross-quantified proteins; Pearson R=0.50, p=1.0×10^-11^, considering proteins significantly affected in both dataset (adj. p < 0.1); Figure S1C). These data indicate that a distinct aging signature is detectable in the liver proteome of NMRs, and that some of the changes in protein abundance already manifest in middle aged (>10 yo) animals.

**Figure 3.**
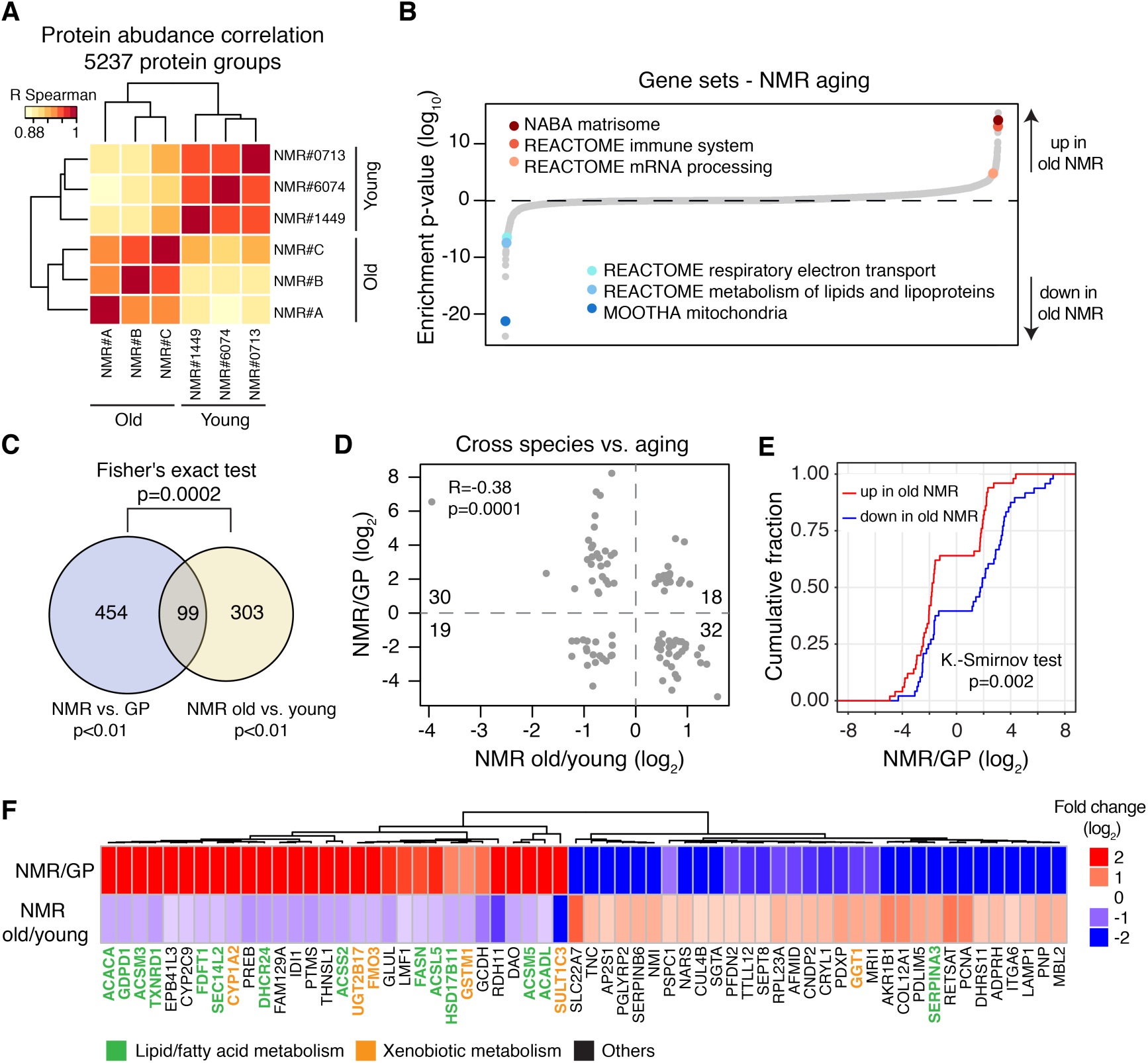
The impact of aging on the NMR liver proteome. **(A)** Livers from 3 young (2.7-3.8 yo) and 3 old (>20 yo) NMRs were compared by Tandem Mass Tags (TMT) based quantitative mass spectrometry. Hierarchical clustering based on the correlation between proteome profiles based on 5237 protein groups cross-quantified between the two age groups (Table S4). **(B)** Gene set enrichment analysis. Gene sets are plotted according to the log_10_ value of the calculated enrichment score. Positive and negative values are used for gene sets showing higher and lower abundance in old NMRs, respectively. Selected significantly affected gene sets (q<0.05) are highlighted. The complete list of enriched gene sets is available in Table S5. **(C)** Overlap between proteins differentially expressed in NMR vs. GP and affected by aging in NMR. **(D)** Comparison between cross-species and aging-related fold changes for the 99 proteins significantly different in both comparisons. **(E)** Cumulative distributions of significant NMR vs. GP fold differences for the 99 proteins also significantly up-(red) or down-(blue) regulated in aging. **(F)** The 62 proteins with significant but opposite fold changes in both comparisons (Table S6). Proteins involved in lipid/fatty acid metabolism and xenobiotic metabolism are highlighted in green and orange, respectively. Related to Figure S1 and Tables S4, S5 and S6.

In order to gain insight into molecular networks affected by aging, we performed Gene Ontology analysis, based on 5237 protein groups quantified by at least 2 unique peptides between young and old NMRs (Table S4). The analysis revealed biological processes that are typically affected by the aging process (Figure 3B). These include increased inflammation and immune response-related proteins [23], and accumulation of extracellular matrix proteins [32]. Interestingly, we found a statistically significant overlap between proteins differentially abundant in NMR vs. GP, and proteins whose abundance is affected by NMR aging (99 proteins, p=2×10^-4^, Fisher’s exact test, Figure 3C). Additionally, we observed a negative correlation between protein fold changes across species and NMR aging (Pearson R=-0.38, p=0.0001; Figure 3D), resulting in a significant difference between cumulative distributions of NMR vs. GP fold changes for proteins up- or down-regulated in NMR aging (p=0.002, Kolmogorov-Smirnov-test; Figure 3E). The directionalities of the differences indicate that proteins with decreasing expression during NMR aging tend to have a higher level in young NMR compared to GP, whereas, proteins with increasing expression during NMR aging tend to start from a lower level in young NMR than in GP. Statistical significance for overlap (p=0.008, Fisher’s exact test), anti-correlation of fold changes (Pearson R=-0.18, p=1.8×10^-8^), and difference between cumulative distributions (p=3.1×10^-12^, Kolmogorov-Smirnov-test) can also be observed from RNA-seq data obtained from the same animals (Figure S1D:F). In particular, we found among the 30 liver proteins up in NMR vs. GP and down during NMR aging 13 linked to lipid or fatty acid and 5 to xenobiotic metabolism (Figure 3F). Our data suggest that this group of proteins is involved in sustaining the longevity of NMRs.

### Aging affects similar pathways in both NMR and human liver

In order to generalize our findings to other species and in particular to human aging, we analyzed the proteome of donor liver samples from 12 individuals aged between 31 and 88 years (Table S1). In this case, we used formalin fixed and paraffin embedded (FFPE) samples and quantitatively compared the proteomes using a new protocol that we recently developed [33]. Principal Component Analysis (PCA) based on 3211 quantified protein groups revealed separation of the proteome profiles based on the age of the donor (Figure 4A). Guided by the PCA analysis, we split the individuals into two groups defined as young (below 50 years of age) and old (above 66 years), and analyzed differential protein expression for 3064 protein groups quantified across all samples (Table S7). Multiple lines of evidence indicate that aging affects similar pathways in both human and NMR liver. First, as in NMR, GO enrichment analysis revealed an age-dependent decline of proteins involved in lipid metabolism and detoxification of xenobiotics, and an increase of proteins related to immune response (Figure 4B and Table S7), and inflammation markers such as RELA/p65 (Figure 4C). Second, enzymes involved in lipid synthesis such as ACACA and DHCR24, which were found to be expressed at higher level in NMR vs. GP and to decline during NMR aging, showed a negative correlation with the age of the donor (Figure 4D and 4E). Enzymes involved in fatty acid beta-oxidation, including ACAA2 and HADHA, also showed a trend of lower abundance in livers from older individuals (Figure S2A). Third, these changes in metabolic enzymes underline a more general reorganization of the liver proteome that is characterized by a significant reduction of both mitochondrial and peroxisomal proteins during aging in both NMR and human (Figure 4F). Fourth, multiple proteins involved in different steps of the xenobiotic metabolism showed similar trends. Cytochrome P450s, a subset of Glutathione S-transferases (GSTs), and UDP-glucuronosyltransferases (UGTs) were in most cases expressed at higher levels in NMR vs. GP and showed an age-dependent decline both in NMR and in human (Figures 4G and S2B). Taken together these data indicate that conserved pathways linked to lipid metabolism and detoxification of xenobiotics are affected in both NMR and humans during aging.

**Figure 4.**
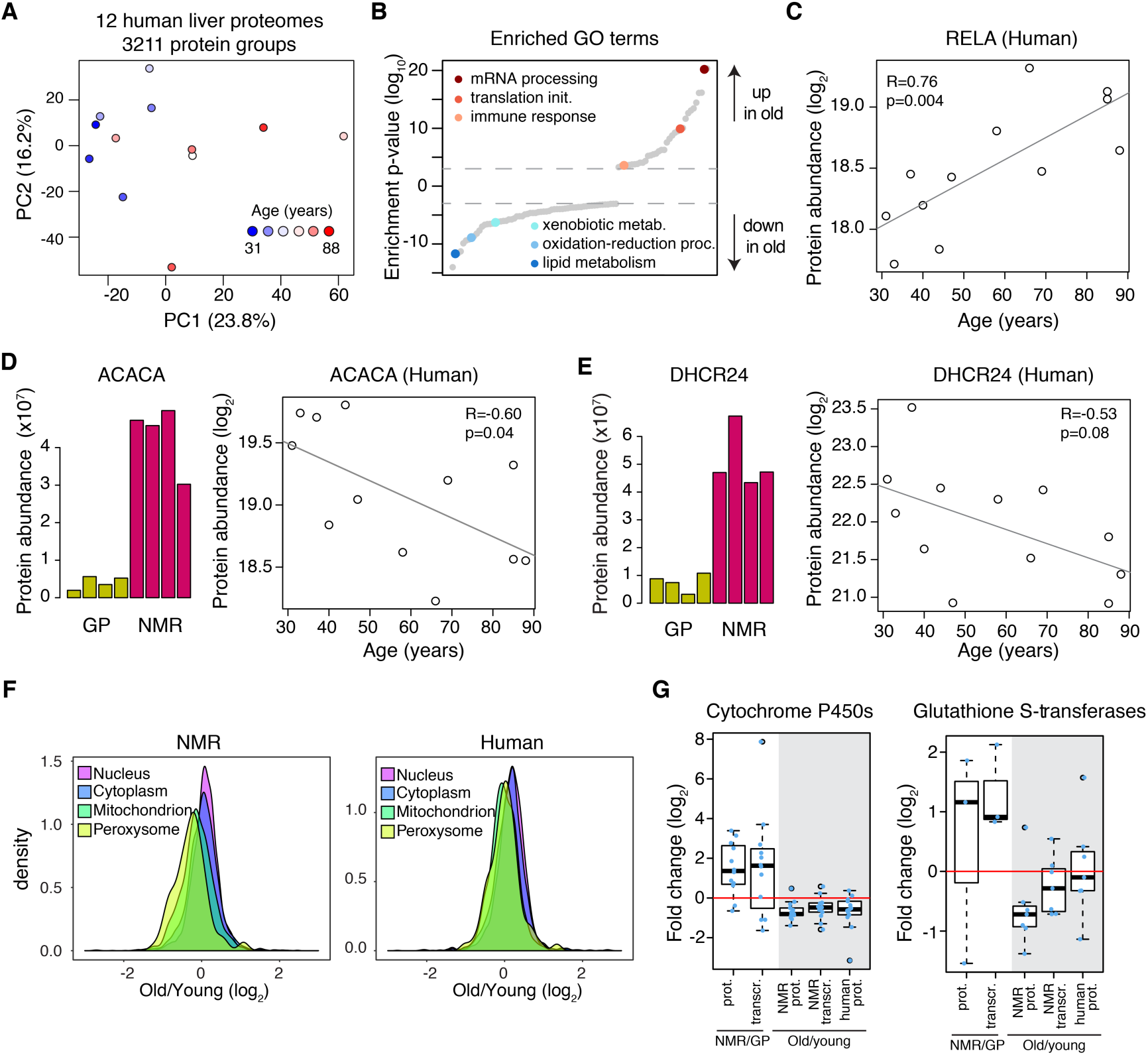
The impact of aging on the human liver proteome and comparison to NMR aging. **(A)** FFPE liver samples from 12 human donors aged 31-88 years old were analysed by Data Independent Acquisition (DIA) quantitative mass spectrometry. Principal Component Analysis (PCA) of the proteome profiles based on 3211 protein groups quantified. **(B)** Gene Ontology (GO) enrichment analysis based on differential protein expression between young (<47 yo, n=6) and old (>66 yo, n=5) donors. One donor aged 58 yo was excluded from differential expression analysis. GO categories are plotted according to the log_10_value of the calculated enrichment p-value. Positive and negative values are used for gene sets showing higher and lower abundance in old individuals, respectively. Selected significantly affected GO terms (p<0.001) are highlighted. The complete list of enriched GO terms is available in Table S7. **(C)** The inflammation marker RELA shows a steady increase of abundance with age. **(D and E)** Selected examples of enzymes involved in lipid metabolism being up regulated in NMR vs. GP and decreasing during aging both in NMRs and humans. **(F)** Mitochondrial and peroxisomal proteins decrease with age in both NMR and human liver. Distributions of fold-changes were calculated separately for proteins assigned to different cellular compartments. Mitochondrial (n=953, p=1.2×10^-37^ Welch two sample t-test for NMR; n=716, p=1.4×10^-21^ Welch two sample t-test for human) and peroxisomal proteins (n=95, p=4.9×10^-6^ Welch two sample t-test for NMR; n= 74, p=8.9×10^-3^ Welch two sample t-test for human) showed a significant lower abundance in old samples as compared to the rest of the quantified proteins. **(G)** Major categories of detoxifying enzymes differentially expressed in NMR vs. GP and affected by aging in both NMR and humans (only significantly affected cases are shown for each group; cut-offs: NMR vs. GP and NMR aging, combined q<0.05; human proteome aging q<0.1). Related to Figure S2 and Table S7.

### The detoxifying enzyme SULT1C3 limits lifespan in C. elegans

In order to demonstrate a functional link between these pathways and longevity, we focused on proteins involved in xenobiotic metabolism and tested their ability to influence lifespan in the nematode *C.elegans.* We chose the phase II enzyme SULT1C3 and its paralog SULT1C2, since these enzymes are among the most prominently expressed in NMR vs. GP, and they progressively decline during aging starting already at 10 years of age (Figure 5A, Figure S1B and Figure S3). The *C. elegans* ortholog of SULTIC2/3 is SSU-1 (coded by *ssu-1*), and it was previously shown to be expressed at high levels in long-lived and stress-resistant dauer larvae (surviving up to 4 months compared to 2-3 weeks lifespan of a reproducing adult). High expression levels of *ssu-1* are thus associated with extended longevity also in worms [34]. The feeding of wild type worms with RNAi against *ssu-1* from L4 larval stage on resulted in significant reduction of their life span (median life span reduction of 13%) suggesting that *ssu-1* is required for normal lifespan of nematodes (Figure 5B). The remarkable stress resistance of dauer larvae is largely due to enhanced expression of pro-survival and detoxification genes mediated by DAF-16/FOXO transcription factor [35,36]. To test whether the FOXO pathway regulates the expression of ssu-1 during adulthood and aging, we performed an epistasis experiment treating *daf-16* deficient nematodes with *ssu-1* RNAi. The lifespan reducing effect of the *ssu-1* RNAi is also observed in *daf-16* mutants (median lifespan reduction of 12%) (Figure 5C), suggesting that the function of *ssu-1* in normal aging is regulated by factors other than DAF-16/FOXO. Taken together these data show that detoxifying enzymes highly expressed in NMR can limit lifespan in a distantly related species.

**Figure 5.**
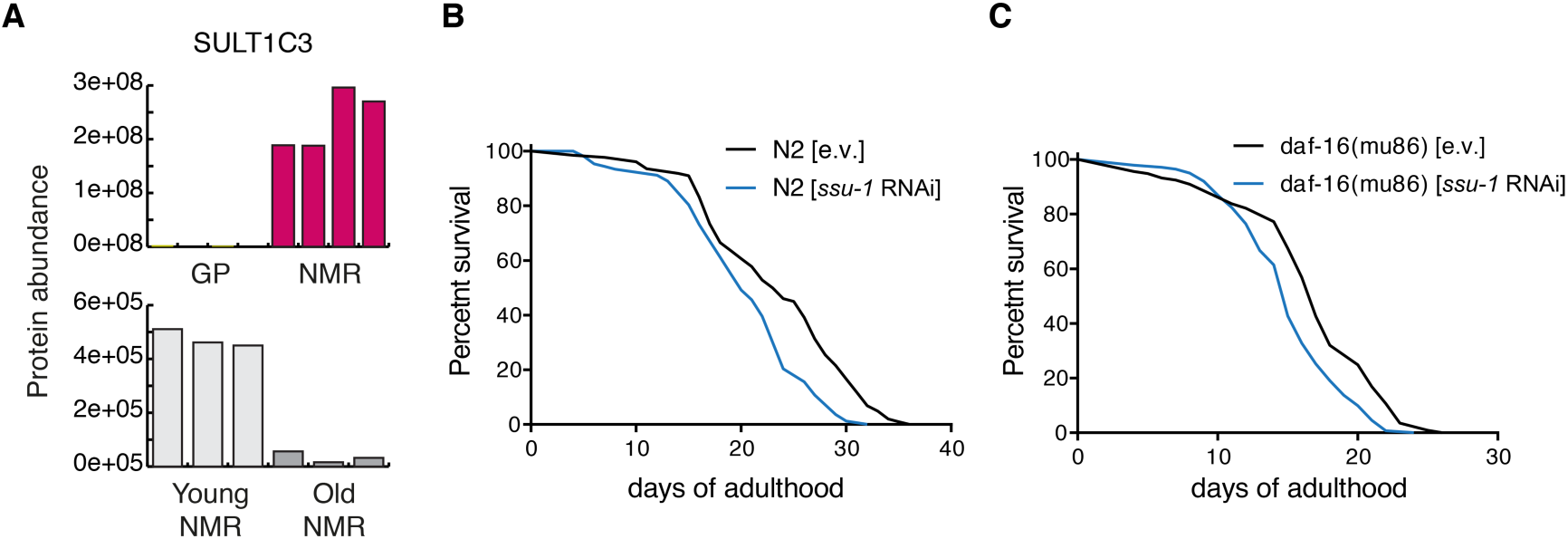
Knock-down of SSU-1 (*ssu-1)* reduces lifespan in wild type and *daf-16(mu86)* mutant *C. elegans*. **(A)** The orthologous mammalian phase II conjugation enzyme SULT1C3 is significantly more abundant in NMR compared to GP, and it declines with aging. **(B)** Wildtype (N2, Bristol strain) and **(C)** *daf-16(mu86)* mutant worms were treated with *ssu-1* and control empty vector (e.v.) RNAi from the L4 (pre-adult) developmental stage. Survival was scored daily; the significance of lifespan shortening was determined by Mantel-Cox Log rank test (p*<*0.0001 in both cases). The N2 experiment was repeated five times and the *daf-16(mu86)* two times with same results. For each repeat, ≥140 animals per condition were used. One representative replicate is shown for each experiment. Related to Figure S3.

## Discussion

Cross-species comparisons based on transcriptional profiling have highlighted pathways that correlate with lifespan [1]. In this work, we decided to concentrate on the NMR, as an outlier of exceptional longevity, and directly relate the identified proteome changes to the ones observed in humans, another of those outliers. In our NMR vs. GP comparative approach, we generally observed a good correlation between transcriptome and proteome differences confirming that a great fraction of adaptation to local ecosystems occurs via changes in gene expression that translate into abundance changes of proteins. However, we have identified changes, particularly among complexes of the mitochondrial respiratory chain, which manifested exclusively at the proteome level. Importantly some of these changes are inline with measureable difference in mitochondrial activity in NMR.

Our cross-species analysis revealed that the liver of NMRs possesses three major characteristics compared to GP: (i) lower rate of mitochondrial respiration, due to reduced level of complex I; (ii) higher reliance on fatty acids for energy production, deriving from increased abundance of enzymes responsible for lipid turnover; and (iii) increased expression of detoxifying enzymes. Although we cannot exclude that some of the observed changes derive from differences in diet between the two compared species, the fact that mitochondrial and oxidative phosphorylation genes were identified to be differentially expressed also in NMRs vs. wild mice [7] supports the peculiarity of NMR liver metabolism among rodents. Importantly, we have also shown a clear impact of aging on the NMR liver proteome that negatively affects the abundance of proteins involved in lipid metabolism and detoxification processes. The same pathways are similarly affected with aging also in mice [37] and humans. This supports the notion of an extremely low, but detectable, rate of aging in NMRs [38].

Two major questions arise from our work: how NMRs have evolved their particular liver metabolism, and how does this contribute to the extreme longevity of these animals? Multiple studies have previously linked the composition of the mitochondrial respiratory chain to lifespan extension in multiple species [31]: altered composition of the respiratory chain has been show to induce a hormetic response that can extend lifespan in *C.elegans* [39]; mild inhibition of complex I leads to increased lifespan in the short-lived fish *N. furzeri* [40]; low abundance of the matrix arm of complex I predicts longevity in mice [41]; fibroblasts isolated from long lived individuals including centenarians show altered mitochondrial activity with lower complex I driven ATP synthesis [42].

Similarly, lipid homeostasis and signaling has been linked to health and longevity [30], and changes in lipid metabolism have been shown to mediate the positive effects of anti-aging dietary interventions. Dietary interventions such as calorie restriction (CR) or fasting influence lipid metabolism [43,44]. Transcriptomic and metabolomic measurements showed that CR promotes fatty acid fueling of mitochondria in liver of mice and it is accompanied by changes in body fat composition [45]. Both fatty acid oxidation and lipid metabolism pathways as well as xenobiotic metabolism are induced by CR in mouse liver via epigenetic reprogramming [46]. Changes in lipid metabolism are mechanistically linked to activation of stress response pathways that mediate enhanced proteostasis [47], and fatty acid oxidation and functional peroxisomes are required to maintain mitochondria network homeostasis and promote longevity in *C. elegans* [48]. In humans, shifts in body composition accompany aging including a decrease of lean mass and accumulation of body fat. Fatty acid oxidation by respiring tissue decreases with age in humans [49,50], however there are discordant reports [51]. A decrease in fatty acid oxidation during aging can lead to adipose tissue accumulation and thus contribute to increased systemic inflammation, a major risk factor for aging-associated disease such as type II diabetes [52]. Our data show that in both NMR and human liver there is a progressive decline of enzymes responsible for fatty acid turnover. These alterations might contribute to changes in energy metabolism that favor the accumulation of adipose tissue and increased inflammation at older age.

From a mechanistic point of view, it is conceivable that adaptation to the particular ecosystem of NMRs has selected for characteristics of energy metabolism that in turn enabled extreme longevity via activation of stress pathways (Figure 6). Among these, the NFE2L2 pathway, which control the expression of many of the detoxifying enzymes that we found increased in NMR vs. GP, was shown to have enhanced activity in NMR [11]. The activity of the same pathways tend to decline during aging, as shown here by the decline of their target genes in both NMR and humans, and in different systems [53,54]. It is therefore tempting to speculate that their higher basal activity in the NMR might contribute to its enhanced stress resistance, and ultimately delay the aging process. In line with this hypothesis, genes encoding for both, respiratory electron transport chain and response to oxidative stress have been shown to be under positive selection in the NMR [6]. Additionally, similar molecular networks (lipid metabolism and oxidative stress pathways) are involved in the social status transition from worker to breeder in NMR [55], suggesting common evolutionary constrains and molecular mechanism underlying longevity and eusociality, i.e. reproductive animals’ lifelong fecundity coupled with extraordinary life- and healthspan in the NMR [56] and even extended lifespan in other mole-rats [57,58] and longevity. Further work is required to elucidate in detail which aspects of liver metabolism are sufficient to promote lifespan, and what is the molecular basis mediating positive systemic effects that support organism health by delaying aging in NMR.

**Figure 6.**
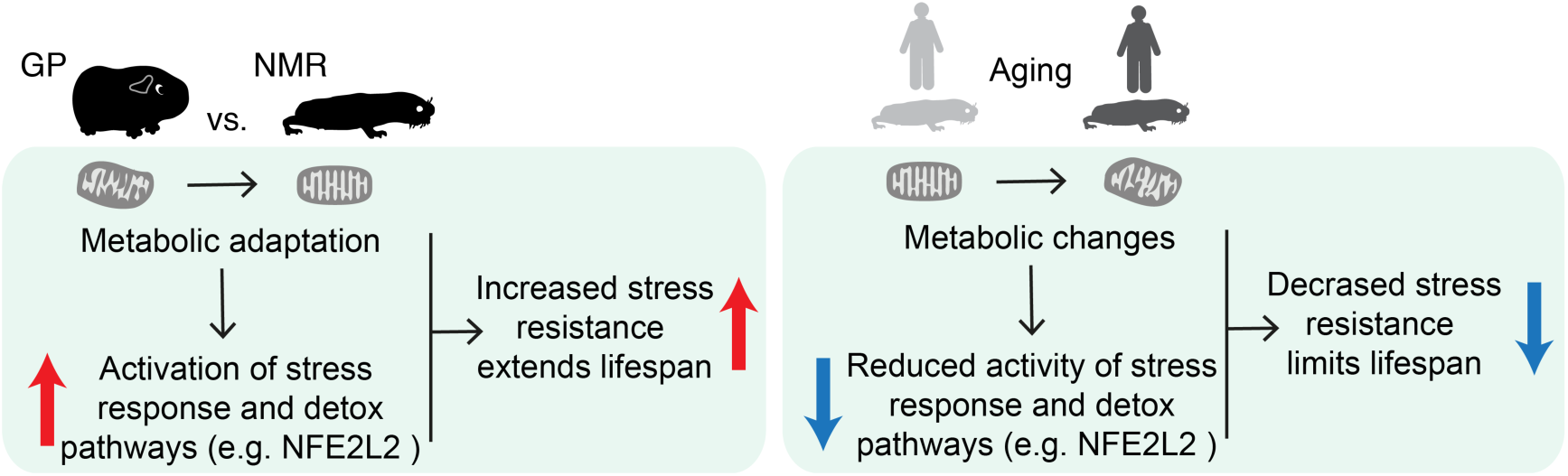
Proposed model linking metabolic changes to stress response/detoxifying pathways in NMR and during aging.

## Materials and methods

### Samples

Young and middle-aged NMR and GP liver tissue samples were obtained at the Leibniz Institute for Zoo and Wildlife Research, IZW (Berlin, Germany). Old NMR liver tissue samples were obtained from the Stockholm Zoo Skansen. Sampling and animal procedures were approved by the local ethics committee of the “Landesamt für Gesundheit und Soziales”, Berlin, Germany (reference numbers: #ZH 156, G02217/12, and T 0073/15) and were compliant with the national and institutional animal care guidelines. NMRs were kept inside artificial burrow systems, consisting of acrylic glass containers interconnected with tubes. Except during cleaning and management procedures, animals were kept in complete darkness and supplied daily *ad libitum* with fresh vegetables. Temperature and humidity were kept stable at 27.0±2.0°C and 85.0±5.0%, respectively. GPs (breed: Dunkin Hartley HsdDhl:DH, Harlan Laboratories, AN Venray, Netherlands) were kept pairwise in standardized GP cages with a 12h light-dark cycle. They were fed commercial pellets and fresh vegetables daily; hay and vitamin C enriched water were provided *ad libitum*. Temperature and humidity ranged between 18.0±2.0°C and 45.0±5.0%, respectively. For tissue collection, animals were euthanized by surgical decapitation under general anaesthesia (Isofluran CP, CP-Pharma, Burgdorf, Germany). Tissue samples were fresh frozen and stored in liquid nitrogen before transcriptome or proteome analysis.

Human liver tissue samples were provided by the tissue bank of the National Center for Tumor Diseases (NCT, Heidelberg, Germany) in accordance with the regulations of the tissue bank and the approval of the ethics committee of Heidelberg University. These samples were obtained in a transplant setting to check the quality of the donor liver by histology before implantation. The age of the donors ranges from 31-88. The tissue samples were formalin fixed, paraffin-embedded (FFPE) and slides were stained with hematoxylin and eosin (H&E). These full-section H&E slides were re-evaluated by a trained pathologist (SS) confirming that each of the samples used for proteomic analyses did not show any overt pathomorphological changes (e.g. necrosis or significant inflammatory or fatty changes).

### RNA sequencing analysis

#### Reference transcripts

Reference transcripts for NMR are based on recently published de novo transcriptome assembly [59]. Reference transcripts for GP were obtained by *de novo* transcriptome assembly of ten different tissue samples as described in [55], using the human transcriptome as a reference for gene symbol assignment. Both transcript sets were mapped to the corresponding genomes (NMR UCSC hetgla2, GP UCSC cavpor3) in two steps: BLAT was used to identify the locus and then SPLIGN (v1.39.8) was applied to splice align the transcript sequence within BLAT locus.

#### Transcript quantification

For NMR age comparison RNA-seq data were aligned to the reference genome utilizing STAR (v2.4.1d) with a maximum mismatch of 6% and a minimum aligned length of 90%. Reads mapped to multiple loci were discarded. Gene expression was quantified using HTSEQ (v0.6.1p1) based on the aligned reference transcripts. For cross-species comparison (NMR vs. GP) orthologous transcribed regions were determined using PosiGene [60] with parameter ‘prank=0 max_anchor_gaps_hard=100 rs=NMR’. RNA-seq data were aligned to corresponding orthologous transcribed regions in NMR and GP reference transcripts using bowtie2 (2.2.9) with the parameter ‘--very-sensitive-local’. DESeq2 (v1.6.3) was used to identify DEGs after correction of p-values using Benjamini Hochberg (FDR, denoted as ‘q’) for NMR aging and cross-species comparison.

### Sample preparation for mass spectrometry

#### NMR and GP fresh frozen liver samples for label free cross species comparison

Frozen tissue samples of NMR and GP livers (between 20 and 40 mg, Table S1) were collected into Precellys Lysing kit tubes (Keramik-kit 1.4/2.8 mm, 2 mL (CKM)) containing 200 μL of protein solubilization buffer (80 μM Tris pH 8.0, 80 μM DTT and 4% SDS) and processed directly. Samples were homogenized in the Precellys 24 homogenizer (Bertin Instruments, France) at 5000 rpm for 30 seconds at 4 °C. Samples were then spun down and the supernatant transferred to a 1.5 mL Eppendorf tube. Samples were sonicated using a Bioruptor Plus (Diagenode) for 7.5 minutes (5 cycles: 1 min on, 30 sec off, 20 °C) using the high setting, and then boiled for 10 min at 95°C. A second round of sonication (as before) followed the boiling. Samples were spun down at 20800x *g* for 5 minutes and the lysate supernatant transferred to fresh tubes. Protein concentration was determined by BCA assay (Pierce) using standard protocol and adjusted to 10 µg/µL using solubilization buffer. 5 µL of tissue lysate, corresponding to 50 µg protein, was taken for preparation for MS. Cysteine residues were alkylated by adding 1 µL of, 200 mM iodoacetamide to a final concentration of 15 mM (incubated for 30 min at room temperature in the dark). Reaction was quenched by addition of 1 μL of 200 mM DTT. Sample clean-up proceeded following a modified SP3 protocol. Sera-Mag Speed Beads (#45152105050250 and #65152105050250, Thermo Scientific) were mixed 1:1, rinsed with water and stored as a 40 μg/μL stock solution in 4°C, as described in [61]. 4 μL of beads stock was added to the reaction tube and mixed by pipetting then 11 µL acetonitrile containing 5% (v/v) formic acid was added. Samples were incubated for 8 minutes at room temperature to allow protein bindings to the beads. Next, tubes were placed on the magnetic rack. Supernatant was removed and discarded. Beads were washed twice with 180 μL of 70% (v/v) ethanol and once with 180 μL of 100% acetonitrile. After removal of acetonitrile beads were air-dried for 60 sec and then resuspended in 7 μL of digestion buffer (6 μL 4 M urea in 100 mM ammonium bicarbonate and 1 μL of 1 μg/μL of LysC (Wako)). Samples were sonicated for 5 min in water bath, incubated for 5 min at 37°C and then mixed by pipetting. Digestion was allowed to proceed for 4 h at 37°C. After the first step of digestion, beads were resuspended by pipetting, urea was diluted to the final concentration of 1.5 M and 1 μL of 1 μg/μL of sequencing grade trypsin (Promega) was added to samples. Digestion was performed overnight at 37°C. After digestion, beads were resuspended by pipetting. 100% acetonitrile was added to the final concentration of 95% (v/v) and samples were incubated for 8 min at room temperature. Tubes were placed on the magnetic rack and washed twice with 100% acetonitrile. Supernatant was removed and beads air-dried and reconstituted in 20 μL of 2% DMSO followed by 5 min of sonication in the water bath. Samples were resuspended by pipetting and placed on the magnetic rack. Supernatant containing peptides was transferred to a fresh tube and acidified with 2 μL of 1% (v/v) formic acid prior to pre-fractionation by high pH reverse phase chromatography. Offline high pH reverse phase fractionation was performed using an Agilent 1260 Infinity HPLC System equipped with a binary pump, degasser, variable wavelength UV detector (set to 220 and 254 nm), peltier-cooled autosampler (set at 10°C) and a fraction collector. The column was a Waters XBridge C18 column (3.5 µm, 100 x 1.0 mm, Waters) with a Gemini C18, 4 x 2.0 mm SecurityGuard (Phenomenex) cartridge as a guard column. The solvent system consisted of 20 mM ammonium formate (pH 10.0) as mobile phase (A) and 100% acetonitrile as mobile phase (B). The separation was accomplished at a mobile phase flow rate of 0.1 mL/min using a linear gradient from 100% A to 35 % B in 61 min. Thirty-four fractions were collected along with the LC separation, which were subsequently pooled into 10 fractions. Pooled fractions were dried in a Speed-Vac and then stored at −80°C until LC-MS/MS analysis.

#### NMR frozen liver samples for TMT-based comparison of young, middle-aged and old samples

For each experimental animal (Table S1), 100 µg protein lysate from the bead-beaten stock of tissue described above were taken up to a final volume of 50 µL with 100 mM HEPES buffer, pH 8.5. 5 µL of 2% SDS was added, prior to biorupting (5 cycles: 1 min on, 30 sec off, 20 °C) at the highest settings. Samples were spun down at 20800x *g* for 1 minute and the lysate supernatant transferred to fresh tubes. Reduction was performed with 2.9 µL DTT (200 mM) for 15 minutes at 45 °C before alkylation with 200 mM IAA (5 µL, 30 minutes, room temperature, in the dark). Proteins were then precipitated with 4 volumes ice cold acetone to 1 volume sample and left overnight at −20 °C. The samples were then centrifuged at 20800x *g* for 30 minutes, 4 °C. After removal of the supernatant, the precipitates were washed twice with 500 µL 80% (v/v) acetone (ice cold). After each wash step, the samples were vortexed, then centrifuged again for 2 minutes at 4°C. The pellets were then allowed to air-dry before being dissolved in digestion buffer (50 µL, 3M urea in 0,1M HEPES, pH 8; 1 µg LysC) and incubated for 4 h at 37 °C with shaking at 600 rpm. Then the samples were diluted 1:1 with milliQ water (to reach 1.5M urea) and were incubated with 1 µg trypsin for 16 h at 37 °C. The digests were then acidified with 10% trifluoroacetic acid and then desalted with Waters Oasis® HLB µElution Plate 30µm in the presence of a slow vacuum. In this process, the columns were conditioned with 3×100 µL solvent B (80% (v/v) acetonitrile; 0.05% (v/v) formic acid) and equilibrated with 3x 100 µL solvent A (0.05% (v/v) formic acid in milliQ water). The samples were loaded, washed 3 times with 100 µL solvent A, and then eluted into PCR tubes with 50 µL solvent B. The eluates were dried down with the speed vacuum centrifuge and dissolved in 200 mM HEPES buffer, pH 8.5 for TMT labeling. 25 µg peptides were taken for each labeling reaction at 1 µg/µL concentration. TMT-6plex reagents for old vs. young comparison (TMT-10plex for middle-aged vs. young comparison) (Thermo Scientific) were reconstituted in 41 µL 100% anhydrous DMSO. TMT labeling was performed by addition of 2.5 μL of the TMT reagent. After 30 minutes of incubation at room temperature, with shaking at 600 rpm in a thermomixer (Eppendorf) a second portion of TMT reagent (2.5 μL) was added and incubated for another 30 minutes. The reaction was quenched with 1 μL of 20 mM lysine in 100 mM ammonium bicarbonate. After checking labeling efficiency, samples were pooled (48 µg total), cleaned once again with Oasis and subjected to high pH fractionation prior to MS analysis. For middle-aged vs. young comparison, two additional samples were generated by pooling the young and middle-aged samples separately and used to fill the two remaining TMT channels. Offline high pH reverse phase fractionation was performed as described above with the following modifications for TMT labeled samples: (i) the separation was accomplished at a mobile phase flow rate of 0.1 mL/min using a non-linear gradient from 95% A to 40 % B in 91 min; (ii) 48 fractions were collected along with the LC separation that were subsequently pooled into 16 fractions.

#### Human FFPE liver samples

The specimens were cut on a microtome into 5 μm thick sections, mounted on glass slides and processed with a modified version of the protocol described in [33]. Slides were deparaffinized in xylene for 2x 5 minutes, rehydrated in 100% ethanol for 2x 5 minutes, and then washed in 96% (v/v), 70% (v/v), 50% (v/v) ethanol and milliQ water for 1x 5 minutes each. Region of interest were gently scraped using a scalpel and transferred to a PCR tube containing 100 μL of protein solubilization buffer (80 μM Tris pH 8.0, 80 μM DTT and 4% SDS) and processed directly. Samples were sonicated using a Bioruptor Plus (Diagenode) for 25.2 min (15 cycles: 1 min on, 30 sec off) at 20°C using the high setting, and then boiled for 1h at 99°C. Sonication followed by boiling was performed twice. Cysteine residues were alkylated by adding 200 mM iodoacetamide to a final concentration of 15 mM (incubated for 30 min at room temperature in the dark). Reaction was quenched by addition of 10 μL of 200 mM DTT. Protein were then acetone precipitated, digested and desalted as described above for NMR samples (aging comparison), with the exceptions that 0.5 µg of both LysC and trypsin were used instead of 1 µg to accommodate for the lower amount of protein extract employed, and no TMT labeling was performed.

### Mass spectrometry data acquisition

#### Label free analysis of NMR and GP liver samples

For label free experiments, each fraction from the 4 GP and 4 NMR samples, separated by high pH, were resuspended in 10 µL reconstitution buffer (5% (v/v) acetonitrile, 0.1% (v/v) TFA in water) and 8 µL were injected. Peptides were separated using the nanoAcquity UPLC system (Waters) fitted with a trapping (nanoAcquity Symmetry C18, 5 µm, 180 µm x 20 mm) and an analytical column (nanoAcquity BEH C18, 2.5 µm, 75 µm x 250 mm). The outlet of the analytical column was coupled directly to an Orbitrap Fusion Lumos (Thermo Fisher Scientific) using the Proxeon nanospray source. Solvent A was water, 0.1% (v/v) formic acid and solvent B was acetonitrile, 0.1% (v/v) formic acid. The samples were loaded with a constant flow of solvent A at 5 µL/min, onto the trapping column. Trapping time was 6 min. Peptides were eluted via the analytical column at a constant flow of 0.3 µL/min, at 40°C. During the elution step, the percentage of solvent B increased in a linear fashion from 5% to 7% in 10 minutes, then from 7% B to 30% B in a further 105 min and to 45% B by 130 min. The peptides were introduced into the mass spectrometer via a Pico-Tip Emitter 360 µm OD x 20 µm ID; 10 µm tip (New Objective) and a spray voltage of 2.2kV was applied. The capillary temperature was set at 300°C. Full scan MS spectra with mass range 375-1500 *m/z* were acquired in profile mode in the Orbitrap with resolution of 120000 FWHM using the quad isolation. A first batch of samples (NMR: F1-6074, M1-1449; GP: #18, #19) was acquired with the following settings. The RF on the ion funnel was set to 60%. The filling time was set at maximum of 100 ms with an AGC target of 4 x 10^5^ ions and 1 microscan. The peptide monoisotopic precursor selection was enabled along with relaxed restrictions if too few precursors were found. The most intense ions (instrument operated for a 3 second cycle time) from the full scan MS were selected for MS2, using quadrupole isolation and a window of 1.6 Da. An intensity threshold of 5 x10^3^ ions was applied. HCD was performed with collision energy of 35%. A maximum fill time of 30 ms with an AGC target of 1 x 10^4^ for each precursor ion was set. MS2 data were acquired in centroid in the ion trap, in Rapid scan mode, with fixed first mass of 120 *m/z*. The dynamic exclusion list was with a maximum retention period of 60 sec and relative mass window of 10 ppm. In order to improve the mass accuracy, internal lock mass correction using a background ion (*m/z* 445.12003) was applied. For data acquisition and processing of the raw data Xcalibur 4.0 (Thermo Scientific) and Tune version 2.0 were employed. As a consequence of method optimization, the following parameters were changed for a second batch of samples (NMR: #0713, #4925; GP: #23, #28): RF on the ion funnel was set to 40%, AGC target to 2 x 10^5^, quadrupole isolation window to 1.4 Da, HCD collision energy to 30%, fill time to 300 ms, AGC target to 2 x 10^3^, and the instrument was set to inject ions for all available parallelizable time. Since the two batches of samples were block randomized (i.e. both contained the same number of NMR and GP samples), the usage of two different methods did not influence the outcome of our comparison, as shown by the expected clustering of the samples according to the species of origin (Figure 1B).

#### TMT analysis of NMR young, middle-aged and old samples

For TMT experiments, fractions were resuspended in 10 µL reconstitution buffer (5% (v/v) acetonitrile, 0.1% (v/v) TFA in water) and 3.5 µL were injected. Peptides were analyzed using the same LC-MS/MS setup described above with the following modifications. Peptides were eluted using a linear gradient from 5% to 7% in 10 minutes, then from 7% B to 30% B in a further 105 min and to 45% B by 130 min. Full scan MS spectra with mass range 375-1500 *m/z* were acquired in profile mode in the Orbitrap with resolution of 60000 FWHM using the quad isolation. The RF on the ion funnel was set to 40%. The filling time was set at maximum of 100 ms with an AGC target of 4 x 10^5^ ions and 1 microscan. The peptide monoisotopic precursor selection was enabled along with relaxed restrictions if too few precursors were found. The most intense ions (instrument operated for a 3 second cycle time) from the full scan MS were selected for MS2, using quadrupole isolation and a window of 1 Da. HCD was performed with collision energy of 35%. A maximum fill time of 50 ms for each precursor ion was set. MS2 data were acquired with fixed first mass of 120 *m/z*. The dynamic exclusion list was with a maximum retention period of 60 sec and relative mass window of 10 ppm. For the MS3, the precursor selection window was set to the range 400-2000 m/z, with an exclude width of 18 *m/z* (high) and 5 *m/z* (low). The most intense fragments from the MS2 experiment were co-isolated (using Synchronus Precursor Selection = 8) and fragmented using HCD (65%). MS3 spectra were acquired in the Orbitrap over the mass range 100-1000 *m/z* and resolution set to 30000 FWMH. The maximum injection time was set to 105 ms and the instrument was set not to inject ions for all available parallelizable time.

#### Data Independent Acquisition (DIA) for human FFPE samples

Peptides were spiked with retention time HRM kit (Biognosys AG), and analyzed using the same LC-MS/MS setup described above with the following modifications. Approx. 1 µg for Data Dependent Acquisition (DDA) and 3 µg for DIA analysis were loaded. Peptides were eluted via a non-linear gradient from 0 % to 40 % in 120 minutes. Total runtime was 145 minutes, including clean-up and column re-equilibration. The RF lens was set to 30%. For spectral library generation, a pooled sample was generated by mixing equal portion of each sample, injected 6 times, and measured in DDA mode. The conditions for DDA data acquisition were as follows: Full scan MS spectra with mass range 350-1650 *m/z* were acquired in profile mode in the Orbitrap with resolution of 60000 FWHM. The filling time was set at maximum of 50 ms with limitation of 2 x 10^5^ ions. The “Top Speed” method was employed to take the maximum number of precursor ions (with an intensity threshold of 5 x 10^4^) from the full scan MS for fragmentation (using HCD collision energy, 30%) and quadrupole isolation (1.4 Da window) and measurement in the Orbitrap (resolution 15000 FWHM, fixed first mass 120 *m/z*), with a cycle time of 3 seconds. The MIPS (monoisotopic precursor selection) peptide algorithm was employed but with relaxed restrictions when too few precursors meeting the criteria were found. The fragmentation was performed after accumulation of 2 x 10^5^ions or after filling time of 22 ms for each precursor ion (whichever occurred first). MS/MS data were acquired in centroid mode. Only multiply charged (2^+^ −7^+^) precursor ions were selected for MS/MS. Dynamic exclusion was employed with maximum retention period of 15s and relative mass window of 10 ppm. Isotopes were excluded. For data acquisition and processing Tune version 2.1 was employed.

For the DIA data acquisition the same gradient conditions were applied to the LC as for the DDA and the MS conditions were varied as follows: Full scan MS spectra with mass range 350-1650 *m/z* were acquired in profile mode in the Orbitrap with resolution of 120000 FWHM. The filling time was set at maximum of 20 ms with limitation of 5 x 10^5^ ions. DIA scans were acquired with 34 mass window segments of differing widths across the MS1 mass range with a cycle time of 3 seconds. HCD fragmentation (30% collision energy) was applied and MS/MS spectra were acquired in the Orbitrap with a resolution of 30000 FWHM over the mass range 200-2000 *m/z* after accumulation of 2 x 10^5^ ions or after filling time of 70 ms (whichever occurred first). Ions were injected for all available parallelizable time. Data were acquired in profile mode.

### Mass spectrometry data analysis

#### Label free cross-species comparison of NMR and GP liver samples

The Andromeda search engine [62], part of MaxQuant (version 1.5.3.28) [63] was used to search the data. The data for GP and NMR were searched separately against translated species-specific reference transcripts (see RNA sequencing analysis). Database with a list of common contaminants were appended in both cases. The data were searched with the following modifications: Carbamidomethyl (C) (Fixed), and Oxidation (M) and Acetyl (Protein N-term) (Variable). The mass error tolerance for the full scan MS spectra was set at 20 ppm and for the MS/MS spectra at 0.5 Da. A maximum of 2 missed cleavages were allowed. Peptide and protein level 1% FDR were applied using a target-decoy strategy [64]. iBAQ (label free quantification) values from the MaxQuant output were used to perform cross-species differential protein expression analysis using scripts written in R (v3.4.1). After removal of reverse and contaminant hits, only protein groups quantified by at least two unique peptides were retained. Common human gene symbols were used to combine iBAQ values for NMR and GP samples. Only protein groups quantified in at least two animals per group were retained when comparing protein abundances between NMR and GP. To reduce technical variation, data were log_2_ transformed and quantile-normalized using the preprocessCore library. Protein differential expression was evaluated using the limma package [65]. Differences in protein abundances were statistically determined using the Student’s t test moderated by the empirical Bayes method. P values were adjusted for multiple testing using the Benjamini-Hochberg method [66] (Table S2).

#### TMT-based analysis of young, middle-aged and old NMR livers

TMT data were processed using Proteome Discoverer v2.0 (Thermo Fisher Scientific). Data were searched against the NMR fasta database using Mascot v2.5.1 (Matrix Science) with the following settings: Enzyme was set to trypsin, with up to 1 missed cleavage. MS1 mass tolerance was set to 10 ppm and MS2 to 0.5 Da. Carbamidomethyl cysteine was set as a fixed modification and oxidation of Methionine as variable. Other modifications included the TMT-6plex modification from the quan method used. The quan method was set for reporter ions quantification with HCD and MS3 (mass tolerance, 20 ppm). The false discovery rate for peptide-spectrum matches (PSMs) was set to 0.01 using Percolator [67]. Reporter ion intensity values for the filtered PSMs were exported and processed using in-house written R scripts to remove common contaminants and decoy hits Additionally only PSMs having reporter ion intensities above 1 x 10^3^ in all the relevant TMT channels were retained for quantitative analysis. Only protein groups quantified by at least two unique peptides were analyzed for differential expression. Data were analysed using the MSnbase package [68]. Reporter ion intensities were log_2_-transformed and normalized using the vsn package [69]. Peptide-level data were summarized into their respective protein groups by taking the median value. Differential protein expression was assessed using the limma package, as described above (Table S4).

#### DIA analysis of FFPE human samples

For library creation, the DDA data was searched using the Andromeda search engine built in MaxQuant (version 1.5.3.28). The data were searched against a human database (Swiss-Prot entries of the Uniprot KB database release 2016_01, 20198 entries) with a list of common contaminants appended, as well as the HRM peptide sequences. The data were searched with the following modifications: Carbamidomethyl (C) (Fixed) and Oxidation (M)/ Acetyl (Protein N-term) (Variable). The mass error tolerance for the full scan MS and MS/MS spectra was set at 20 ppm. A maximum of 1 missed cleavage was allowed. The identifications were filtered to satisfy FDR of 1 % on peptide and protein level. A spectral library was created from the MaxQuant output of the DDA runs combined using Spectronaut (version 10, Biognosys AG). This library contained 34014 precursors, corresponding to 3295 protein groups using Spectronaut protein inference. DIA data were then uploaded and searched against this spectral library. Precursor matching, protein inference and quantification was performed in Spectronaut using default settings [70]. Differential protein expression was evaluated using a pairwise t-test performed at the precursor level followed by multiple testing correction according to [71]. The data (candidate table, Table S7) was exported from Spectronaut and used for further data analyses (see below).

### Data analysis

For integrated analysis of RNA-seq and proteomic data, transcripts and protein groups were matched using the corresponding gene symbol, P values were combined using the Fisher method, followed by correction for multiple testing using the Benjamini-Hochberg method [66]. Gene set enrichments (Figures 2A and 3B) were performed with the R package *gage* [72] using gene set definitions from the Molecular Signatures Database (MSigDB, C2 v5.1) [73]. Gene Ontology enrichment analysis (Figure 4B) was performed the list of quantified proteins that were ranked according to the level of differential expression (fold change) using GOrilla (Eden et al., 2009) followed by GO term redundancy reduction performed by REViGO [74] using default settings.

### Measurements of mitochondrial activity

Mitochondrial respiration was measured in homogenized liver tissue samples of NMRs and mice by means of high-resolution respirometry using the Oroboros^®^ Oxygraph-2K (Oroboros Instruments, Innsbruck, Austria). This device allows for simultaneous recording the O_2_ concentration in two parallel chambers calibrated for 2 ml of respiration medium containing 110 mM D-Sucrose (Sigma 84097), 60 mM K-lactobionate (Aldrich 153516), 0.5 mM ethylene glycol tetra acetic acid (Sigma E4378), 1 g/L bovine serum albumin free from essentially fatty acids (Sigma A 6003), 3 mM MgCl_2_ (Scharlau MA0036), 20 mM taurine (Sigma T0625), 10 mM KH_2_PO_4_ (Merck 104873), 20mM HEPES (Sigma H7523), adjusted to pH 7.1 with KOH and equilibrated with 21% O_2_ at 37°C. Mitochondrial respiration was quantified in terms of oxygen flux (*J*O_2_) calculated as the rate of change of the O_2_ concentration in the chambers normalized for wet tissue volume.

The liver tissue homogenates were generated from 40-50 mg of wet tissue samples suspended in 2 ml of ice-cold respiration medium. Aliquots of the homogenates were added to each oxygraph chamber in order to obtain a final amount of 4 mg of NMR liver tissue or 2 mg of mouse liver tissue per chamber. The different amount of tissue was chosen in order to obtain similar absolute *J*O2-values, i.e. *J*O2-values not normalized per wet weight, in both species. Every sample was measured in duplicates; the mean values from both chambers were used for statistical analysis.

The titration sequence used for the experiments was as follows: 2 mM malate + 1 mM octanoylcarnitine, 5 mM ADP, 10 μM cytochrome c, 10 mM glutamate, 10 mM succinate, 2.5 μM oligomycin, 1 μM carbonyl cyanide p-(trifluoromethoxy)-phenylhydrazone (FCCP), 0.5 μM rotenone, and 5 μM antimycin A. The *J*O_2_-values after addition of octanoyl carnitine, malate, ADP, and cytochrome c allow quantifying fatty acid oxidation. The addition of cytochrome c after ADP is required to test the integrity of the outer mitochondrial membrane. If homogenization steps damaged the mitochondrial membrane, addition of cytochrome c induces an increase of the respiratory values. The maximum oxidative capacity of the mitochondrial respiratory chain in the coupled state (maximum OxPhos) was then determined after the subsequent addition of glutamate and succinate. Further injections of the ATP synthase inhibitor oligomycin and of the uncoupler FCCP allowed obtaining the maximum respiratory activity in the uncoupled state. In the next two steps, complex I and complex III were sequentially inhibited by administration of rotenone and antimycin A respectively. Finally sequentially injecting 2 mM ascorbate and 0.5 mM of the complex IV substrate tetramethyl phenylene diamine (TMPD) in the parallel chambers allowed for selectively quantifying the activity of the cytochrome-c-oxidase (COX). Part of the *J*O_2_ induced by the injection of TMPD is caused by auto-oxidation of this compound. Therefore, inhibiting the COX by 40 µM sodium sulfide allowed to quantify and thus to subtract this auto-oxidation related part from the total *J*O_2_ value under TMPD.

### C. elegans lifespan measurements

HT115 bacteria containing specific RNAi constructs were grown on lysogeny broth agar plates supplemented with ampicillin and tetracycline. Plates were kept at 4°C. Overnight cultures were grown in lysogeny broth media containing ampicillin. RNAi expression was induced by adding 1 mM isopropylthiogalactoside (IPTG) and incubating the cultures at 37°C for 20 minutes before seeding bacteria on NGM agar supplemented with ampicillin and 3 mM IPTG. Synchronized L4 larvae were placed on 60 mm dishes containing RNAi expressing bacteria at a density of 70 worms per plate. Worms were transferred to new plates on a daily basis until adulthood day 6 (AD6) and later transferred to new plates every 3-4 days. The number of dead animals was scored daily. The analysis of the lifespan data including statistics was performed using GraphPad Prism software.

### Data availability

RNA-seq data of GP used in *de novo* assembly were deposited at Sequence Read Archive (SRP104222). RNA-seq data for NMR age and cross-species comparisons were deposited in Gene Expression Omnibus (GSE98744). The corresponding gene annotation for NMR and GP are available as a gff3-file (ftp://genome.leibniz-fli.de/pub/nmr2017/). The mass spectrometry proteomics data have been deposited to the ProteomeXchange Consortium via the PRIDE [75] partner repository with the dataset identifier PXD008720.

## Acknowledgements

We thank staff members from Stockholm Zoo Skansen for providing old NMRs, Veronika Geißler and other members of NCT tissue bank for their support, and Stefan Pietsch for support with bioinformatics analysis. We gratefully acknowledge support from the FLI proteomics, sequencing, and bioinformatics core facilities. The FLI is a member of the Leibniz Association and is financially supported by the Federal Government of Germany and the State of Thuringia.

## Author contributions

IH, EC, SH, MV performed experiments and analysed data; MB, AS, KS, NR analysed data; MP, TH, AO, SS, JMK, KS, EC, ME designed experiments; MP, TH and AO oversaw the project and wrote the manuscript.

## Figures

**Figure S1.**
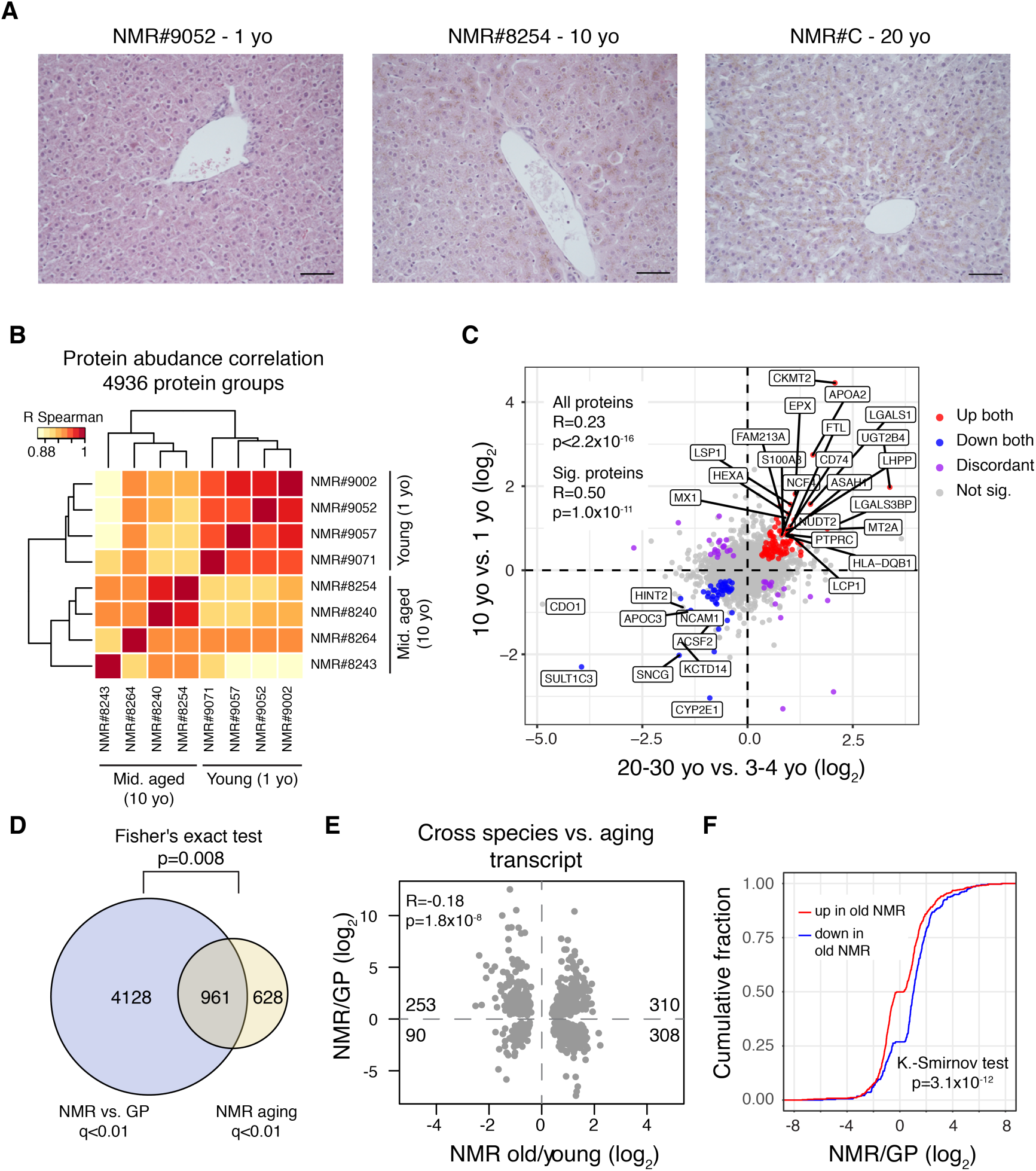
Comparison of liver proteomes between middle-aged and young NMRs, and correlation analysis of transcript level differences in NMR vs. GP and NMR aging. **(A)** Representative micrographs show H&E stained (FFPE) liver tissue sections from NMRs of the indicated age groups. Scale bar = 100 µm**. (B)** Livers from 4 young (1 yo) and 4 middle-aged (>10 yo) NMRs were compared by Tandem Mass Tags (TMT) based quantitative mass spectrometry. Hierarchical clustering based on the correlation between proteome profiles based on 4936 protein groups cross-quantified between the two age groups (Table S4). **(C)** Comparison of protein fold changes calculated between old vs. young (x-axis) and middle-aged vs. young (y-axis) NMRs. Colored dots indicate proteins significant (adj. p<0.1) in both comparisons. The names of selected proteins that show consisted fold changes in the two comparisons are indicated. **(D)** Significant overlap between differentially expressed genes (DEGs) in NMR vs. GP and in aging of NMR. **(E)** Comparison between cross-species and aging-related fold changes for the 875 DEGs significant in both comparison (q<0.01) shows significant negative correlation. **(F)** Cumulative distributions of NMR vs. GP fold changes for the 875 DEGs also significantly up-(red) or down-(blue) regulated in NMR aging. The x-axis was restricted to ±8 for display purpose. Related to Figure 3, Table S4 and Table S6.

**Figure S2.**
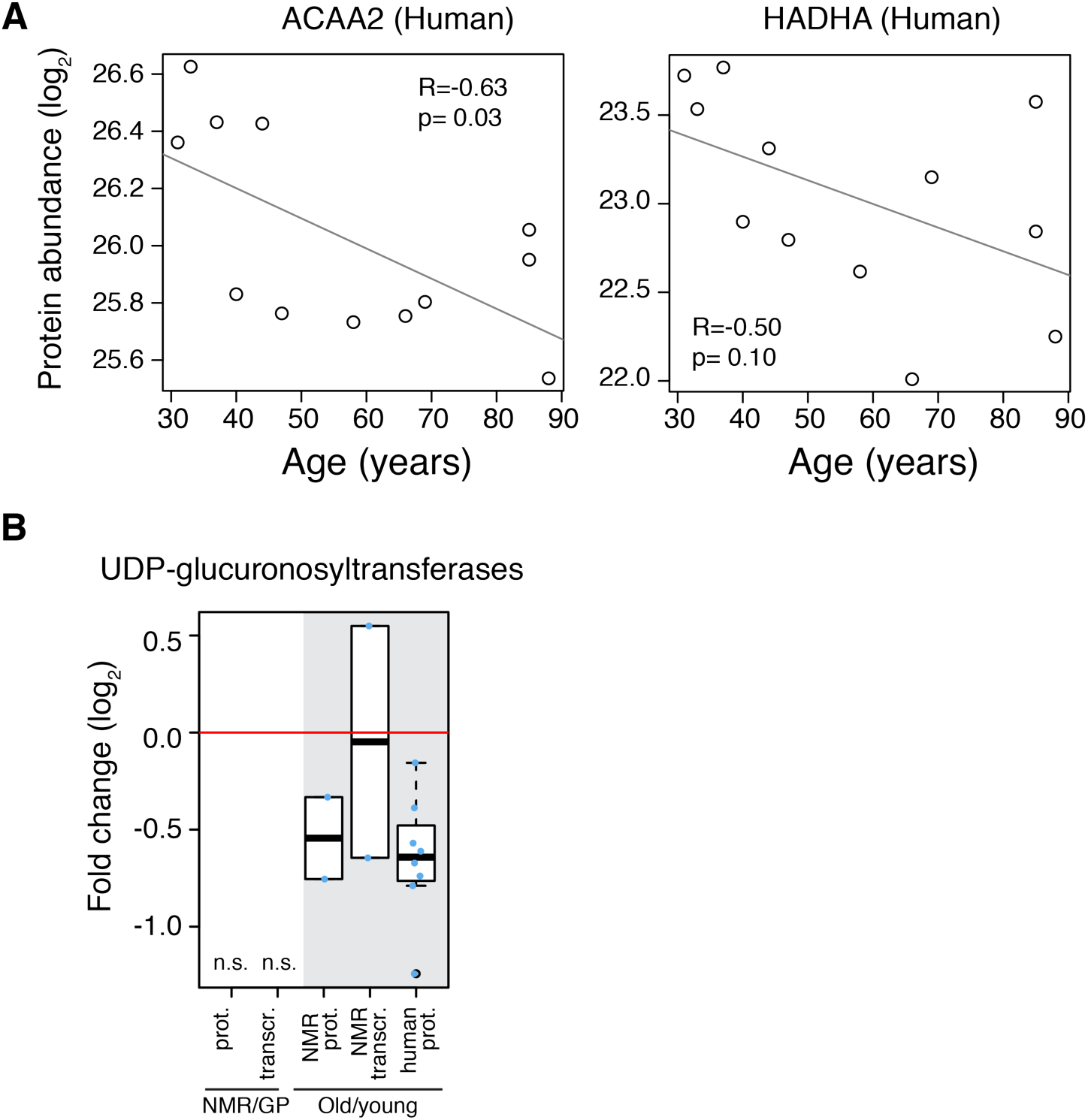
Examples of enzymes involved in fatty acid beta-oxidation and xenobiotic metabolism that decrease during aging in human liver. **(A)** Additional examples of enzymes involved in lipid metabolism decreasing during aging in human liver. **(B)** Detoxifying enzymes decreasing during aging in both NMR and humans. Only significantly affected genes are shown; cut offs: NMR aging, combined q<0.05; human proteome aging q<0.1; n.s. = no significant cases detected. Related to Figure 4.

**Figure S3.**
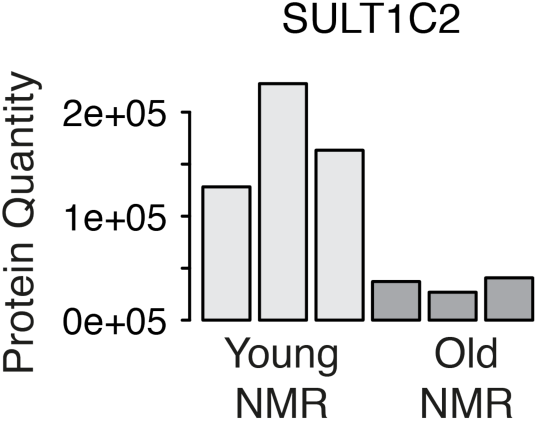
SULT1C2 decreases with aging in NMR. Related to Figure 5.

## Additional files

- **Table S1:** Excel workbook (.xlsx), list of used animals.
- **Table S2:** Excel workbook (.xlsx), NMR vs. GP dataset, contains differential expression analysis for proteomic and transcriptomic data and integrated RNAseq proteome analysis of NMR and GP livers.
- **Table S3:** Excel workbook (.xlsx), NMR vs. GP gene set enrichment, contains significantly affected gene sets from the proteomic analysis.
- **Table S4:** Excel workbook (.xlsx), NMR aging dataset, contains differential expression analysis for proteomic and transcriptomic data and integrated RNAseq proteome analysis of old vs. young NMR livers, and proteomic analysis of middle-aged vs. young NMR livers.
- **Table S5:** Excel workbook (.xlsx), NMR aging gene set enrichment, contains significantly affected gene sets from the proteomic analysis.
- **Table S6:** Excel workbook (.xlsx), comparison of cross-species and NMR aging dataset for proteomic and transcriptomic data.
- **Table S7:** Excel workbook (.xlsx), human aging dataset, contains differential expression analysis for proteomic data for human liver samples from young and old donors.

## References

1. Fushan AA, Turanov AA, Lee S-G, Kim EB, Lobanov A V, Yim SH, et al. Gene expression defines natural changes in mammalian lifespan. Aging Cell. 2015;352–65.

2. Sahm A, Bens M, Szafranski K, Holtze S, Groth M, Goerlach M, et al. Long-lived rodents reveal signatures of positive selection in genes associated with lifespan and eusociality. doi.org. 2017;191999.

3. Lewis KN, Mele J, Hornsby PJ, Buffenstein R. Stress resistance in the naked mole-rat: the bare essentials - a mini-review. Gerontology. 2012;453–62.

4. Skulachev VP, Holtze S, Vyssokikh MY, Bakeeva LE, Skulachev M V., Markov A V., et al. Neoteny, Prolongation of Youth: From Naked Mole Rats to “Naked Apes” (Humans). Physiol. Rev. 2017;699–720.

5. Kim EB, Fang X, Fushan AA, Huang Z, Lobanov A V., Han L, et al. Genome sequencing reveals insights into physiology and longevity of the naked mole rat. Nature. 2011;223– 7.

6. Fang X, Seim I, Huang Z, Gerashchenko M V., Xiong Z, Turanov AA, et al. Adaptations to a Subterranean Environment and Longevity Revealed by the Analysis of Mole Rat Genomes. Cell Rep. 2014;1354–64.

7. Yu C, Li Y, Holmes A, Szafranski K, Faulkes CG, Coen CW, et al. RNA sequencing reveals differential expression of mitochondrial and oxidation reduction genes in the long-lived naked mole-rat when compared to mice. PLoS One. 2011;e26729.

8. Pérez VI, Buffenstein R, Masamsetti V, Leonard S, Salmon AB, Mele J, et al. Protein stability and resistance to oxidative stress are determinants of longevity in the longest-living rodent, the naked mole-rat. Proc. Natl. Acad. Sci. U. S. A. 2009;3059–64.

9. Rodriguez KA, Edrey YH, Osmulski P, Gaczynska M, Buffenstein R. Altered composition of liver proteasome assemblies contributes to enhanced proteasome activity in the exceptionally long-lived naked mole-rat. PLoS One. 2012;e35890.

10. Buffenstein R. Negligible senescence in the longest living rodent, the naked mole-rat: Insights from a successfully aging species [Internet]. J. Comp. Physiol. B Biochem. Syst. Environ. Physiol. 2008. p. 439–45.

11. Lewis KN, Wason E, Edrey YH, Kristan DM, Nevo E, Buffenstein R. Regulation of Nrf2 signaling and longevity in naturally long-lived rodents. Proc. Natl. Acad. Sci. U. S. A. 2015;3722–7.

12. Tian X, Azpurua J, Hine C, Vaidya A, Myakishev-Rempel M, Ablaeva J, et al. High-molecular-mass hyaluronan mediates the cancer resistance of the naked mole rat. Nature. 2013;346–9.

13. Andziak B, O’Connor TP, Qi W, DeWaal EM, Pierce A, Chaudhuri AR, et al. High oxidative damage levels in the longest-living rodent, the naked mole-rat. Aging Cell. 2006;463–71.

14. Holtze S, Eldarov CM, Vays VB, Vangeli IM, Vysokikh MY, Bakeeva LE, et al. Study of age-dependent structural and functional changes of mitochondria in skeletal muscles and heart of naked mole rats (Heterocephalus glaber). Biochem. 2016;1429–37.

15. Lewis KN, Andziak B, Yang T, Buffenstein R. The naked mole-rat response to oxidative stress: just deal with it. Antioxid. Redox Signal. 2013;1388–99.

16. Finkel T. The metabolic regulation of aging. Nat. Med. 2015;1416–23.

17. Fontana L, Partridge L. Promoting Health and Longevity through Diet: From Model Organisms to Humans. Cell. 2015;106–18.

18. Davinelli S, Willcox DC, Scapagnini G. Extending healthy ageing: nutrient sensitive pathway and centenarian population. Immun. Ageing. 2012;9.

19. Sahm A, Bens M, Platzer M, Cellerino A. Parallel evolution of genes controlling mitonuclear balance in short-lived annual fishes. Aging Cell. 2017 Feb 11;

20. Houtkooper RH, Mouchiroud L, Ryu D, Moullan N, Katsyuba E, Knott G, et al. Mitonuclear protein imbalance as a conserved longevity mechanism. Nature. 2013;451–7.

21. Brandt T, Mourier A, Tain LS, Partridge L, Larsson N-G, Kühlbrandt W. Changes of mitochondrial ultrastructure and function during ageing in mice and Drosophila. Elife. 2017;

22. Park TJ, Reznick J, Peterson BL, Blass G, Omerbašić D, Bennett NC, et al. Fructose-driven glycolysis supports anoxia resistance in the naked mole-rat. Science (80-.). 2017;307–11.

23. Ori A, Toyama BH, Harris MS, Bock T, Iskar M, Bork P, et al. Integrated Transcriptome and Proteome Analyses Reveal Organ-Specific Proteome Deterioration in Old Rats. Cell Syst. 2015;224–37.

24. Schwanhäusser B, Busse D, Li N, Dittmar G, Schuchhardt J, Wolf J, et al. Global quantification of mammalian gene expression control. Nature. 2011;337–42.

25. Vogel C, Marcotte EM. Insights into the regulation of protein abundance from proteomic and transcriptomic analyses. Nat. Rev. Genet. 2012;227–32.

26. Nyström T, Yang J, Molin M. Peroxiredoxins, gerontogenes linking aging to genome instability and cancer. Genes Dev. 2012;2001–8.

27. Hanzén S, Vielfort K, Yang J, Roger F, Andersson V, Zamarbide-Forés S, et al. Lifespan Control by Redox-Dependent Recruitment of Chaperones to Misfolded Proteins. Cell. 2016;140–51.

28. Biteau B, Karpac J, Supoyo S, DeGennaro M, Lehmann R, Jasper H. Lifespan extension by preserving proliferative homeostasis in Drosophila. Kim SK, editor. PLoS Genet. 2010;1–15.

29. Erol A. The Functions of PPARs in Aging and Longevity. PPAR Res. 2007;39654.

30. Bustos V, Partridge L. Good Ol’ Fat: Links between Lipid Signaling and Longevity. Trends Biochem. Sci. 2017;

31. Cellerino A, Ori A. What have we learned on aging from omics studies? Semin. Cell Dev. Biol. 2017;

32. Delire B, Lebrun V, Selvais C, Henriet P, Bertrand A, Horsmans Y, et al. Aging enhances liver fibrotic response in mice through hampering extracellular matrix remodeling. Aging (Albany. NY). 2016;98–113.

33. Buczak K, Ori A, Kirkpatrick JM, Holzer K, Dauch D, Roessler S, et al. Spatial tissue proteomics quantifies inter- and intra-tumor heterogeneity in hepatocellular carcinoma. Mol. Cell. Proteomics. 2018;

34. Hattori K, Inoue M, Inoue T, Arai H, Tamura H. A novel sulfotransferase abundantly expressed in the dauer larvae of Caenorhabditis elegans. J. Biochem. 2006;355–62.

35. Fielenbach N, Antebi A. C. elegans dauer formation and the molecular basis of plasticity. Genes Dev. 2008;2149–65.

36. Murphy CT, McCarroll SA, Bargmann CI, Fraser A, Kamath RS, Ahringer J, et al. Genes that act downstream of DAF-16 to influence the lifespan of Caenorhabditis elegans. Nature. 2003;277–83.

37. White RR, Milholland B, MacRae SL, Lin M, Zheng D, Vijg J. Comprehensive transcriptional landscape of aging mouse liver. BMC Genomics. 2015;899.

38. Beltrán-Sánchez H, Finch C. Age is just a number. Elife. 2018;

39. Durieux J, Wolff S, Dillin A. The cell-non-autonomous nature of electron transport chain-mediated longevity. Cell. 2011;79–91.

40. Baumgart M, Priebe S, Groth M, Hartmann N, Menzel U, Pandolfini L, et al. Longitudinal RNA-seq analysis of vertebrate aging identifies mitochondrial complex i as a small-molecule-sensitive modifier of lifespan. Cell Syst. 2016;122–32.

41. Miwa S, Jow H, Baty K, Johnson A, Czapiewski R, Saretzki G, et al. Low abundance of the matrix arm of complex I in mitochondria predicts longevity in mice. Nat. Commun. 2014;3837.

42. Sgarbi G, Matarrese P, Pinti M, Lanzarini C, Ascione B, Gibellini L, et al. Mitochondria hyperfusion and elevated autophagic activity are key mechanisms for cellular bioenergetic preservation in centenarians. Aging (Albany. NY). 2014;296–310.

43. Sengupta S, Peterson TR, Laplante M, Oh S, Sabatini DM. mTORC1 controls fasting-induced ketogenesis and its modulation by ageing. Nature. 2010;1100–4.

44. Luis NM, Wang L, Ortega M, Deng H, Katewa SD, Li PW-L, et al. Intestinal IRE1 Is Required for Increased Triglyceride Metabolism and Longer Lifespan under Dietary Restriction. Cell Rep. 2016;1207–16.

45. Mitchell SJ, Madrigal-Matute J, Scheibye-Knudsen M, Fang E, Aon M, González-Reyes JA, et al. Effects of Sex, Strain, and Energy Intake on Hallmarks of Aging in Mice. Cell Metab. 2016;1093–112.

46. Hahn O, Grönke S, Stubbs TM, Ficz G, Hendrich O, Krueger F, et al. Dietary restriction protects from age-associated DNA methylation and induces epigenetic reprogramming of lipid metabolism. Genome Biol. 2017;1194.

47. Kim HE, Grant AR, Simic MS, Kohnz RA, Nomura DK, Durieux J, et al. Lipid Biosynthesis Coordinates a Mitochondrial-to-Cytosolic Stress Response. Cell. 2016;1539–1552.e16.

48. Weir HJ, Yao P, Huynh FK, Escoubas CC, Goncalves RL, Burkewitz K, et al. Dietary Restriction and AMPK Increase Lifespan via Mitochondrial Network and Peroxisome Remodeling. Cell Metab. 2017;

49. Toth M, Tchernof A. Lipid metabolism in the elderly. Eur. J. Clin. Nutr. 2000;S121–5.

50. Solomon TPJ, Marchetti CM, Krishnan RK, Gonzalez F, Kirwan JP. Effects of aging on basal fat oxidation in obese humans. Metabolism. 2008;1141–7.

51. St-Onge M-P, Gallagher D. Body composition changes with aging: the cause or the result of alterations in metabolic rate and macronutrient oxidation? Nutrition. 2010;152–5.

52. Franceschi C, Campisi J, LR M, J C, JL K, HY C. Chronic Inflammation (Inflammaging) and Its Potential Contribution to Age-Associated Diseases. Journals Gerontol. Ser. A Biol. Sci. Med. Sci. 2014;S4–9.

53. Done AJ, Gage MJ, Nieto NC, Traustadóttir T. Exercise-induced Nrf2-signaling is impaired in aging. Free Radic. Biol. Med. 2016;130–8.

54. Safdar A, deBeer J, Tarnopolsky MA. Dysfunctional Nrf2–Keap1 redox signaling in skeletal muscle of the sedentary old. Free Radic. Biol. Med. 2010;1487–93.

55. Bens M, Szafranski K, Holtze S, Sahm A, Groth M, Kestler HA, et al. Naked mole-rat transcriptome signatures of socially-suppressed sexual maturation and links of reproduction to aging. bioRxiv. 2017;221333.

56. Buffenstein R, Jarvis JUM. The Naked Mole Rat--A New Record for the Oldest Living Rodent. Sci. Aging Knowl. Environ. 2002;7pe–7.

57. Dammann P, Šumbera R, Maßmann C, Scherag A, Burda H. Extended Longevity of Reproductives Appears to be Common in Fukomys Mole-Rats (Rodentia, Bathyergidae). de Polavieja GG, editor. PLoS One. 2011;e18757.

58. Dammann P, Burda H. Sexual activity and reproduction delay ageing in a mammal. Curr. Biol. 2006;R117–8.

59. Bens M, Sahm A, Groth M, Jahn N, Morhart M, Holtze S, et al. FRAMA: from RNA-seq data to annotated mRNA assemblies. BMC Genomics. 2016;54.

60. Sahm A, Bens M, Platzer M, Szafranski K. PosiGene: automated and easy-to-use pipeline for genome-wide detection of positively selected genes. Nucleic Acids Res. 2017;e100–e100.

61. Hughes CS, Foehr S, Garfield DA, Furlong EE, Steinmetz LM, Krijgsveld J. Ultrasensitive proteome analysis using paramagnetic bead technology. Mol. Syst. Biol. 2014;757.

62. Cox J, Neuhauser N, Michalski A, Scheltema RA, Olsen J V., Mann M. Andromeda: A Peptide Search Engine Integrated into the MaxQuant Environment. J. Proteome Res. 2011;1794–805.

63. Cox J, Mann M. MaxQuant enables high peptide identification rates, individualized p.p.b.-range mass accuracies and proteome-wide protein quantification. Nat. Biotechnol. 2008;1367–72.

64. Elias JE, Gygi SP. Target-decoy search strategy for increased confidence in large-scale protein identifications by mass spectrometry. Nat. Methods. 2007;207–14.

65. Ritchie ME, Phipson B, Wu D, Hu Y, Law CW, Shi W, et al. limma powers differential expression analyses for RNA-sequencing and microarray studies. Nucleic Acids Res. 2015;e47–e47.

66. Benjamini Y, Hochberg Y. Controlling the False Discovery Rate: A Practical and Powerful Approach to Multiple Testing [Internet]. J. R. Stat. Soc. Ser. B. 1995. p. 289–300.

67. Brosch M, Yu L, Hubbard T, Choudhary J. Accurate and sensitive peptide identification with Mascot Percolator. J. Proteome Res. 2009;3176–81.

68. Gatto L, Lilley KS. MSnbase-an R/Bioconductor package for isobaric tagged mass spectrometry data visualization, processing and quantitation. Bioinformatics. 2012;288–9.

69. Huber W, von Heydebreck A, Sültmann H, Poustka A, Vingron M. Variance stabilization applied to microarray data calibration and to the quantification of differential expression. Bioinformatics. 2002;S96–104.

70. Bruderer R, Bernhardt OM, Gandhi T, Miladinović SM, Cheng L-Y, Messner S, et al. Extending the limits of quantitative proteome profiling with data-independent acquisition and application to acetaminophen-treated three-dimensional liver microtissues. Mol. Cell. Proteomics. 2015;1400–10.

71. Storey JD. A direct approach to false discovery rates. J. R. Stat. Soc. Ser. B (Statistical Methodol. 2002;479–98.

72. Luo W, Friedman MS, Shedden K, Hankenson KD, Woolf PJ. GAGE: generally applicable gene set enrichment for pathway analysis. BMC Bioinformatics. 2009;161.

73. Subramanian A, Tamayo P, Mootha VK, Mukherjee S, Ebert BL, Gillette MA, et al. Gene set enrichment analysis: a knowledge-based approach for interpreting genome-wide expression profiles. Proc. Natl. Acad. Sci. U. S. A. 2005;15545–50.

74. Supek F, Bošnjak M, Škunca N, Šmuc T. REVIGO summarizes and visualizes long lists of gene ontology terms. PLoS One. 2011;e21800.

75. Vizcaíno JA, Csordas A, Del-Toro N, Dianes JA, Griss J, Lavidas I, et al. 2016 update of the PRIDE database and its related tools. Nucleic Acids Res. 2016;D447–56.

